# Epitope-based vaccine design yields fusion peptide-directed antibodies that neutralize diverse strains of HIV-1

**DOI:** 10.1101/306282

**Authors:** Kai Xu, Priyamvada Acharya, Rui Kong, Cheng Cheng, Gwo-Yu Chuang, Kevin Liu, Mark K. Louder, Sijy O’Dell, Reda Rawi, Mallika Sastry, Chen-Hsiang Shen, Baoshan Zhang, Tongqing Zhou, Mangaiarkarasi Asokan, Robert T. Bailer, Michael Chambers, Xuejun Chen, Chang W. Choi, Venkata P. Dandey, Nicole A. Doria-Rose, Aliaksandr Druz, Edward T. Eng, S. Katie Farney, Kathryn E. Foulds, Hui Geng, Ivelin S. Georgiev, Jason Gorman, Kurt R. Hill, Alexander J. Jafari, Young D. Kwon, Yen-Ting Lai, Thomas Lemmin, Krisha McKee, Tiffany Y. Ohr, Li Ou, Dongjun Peng, Ariana P. Rowshan, Zizhang Sheng, John-Paul Todd, Yaroslav Tsybovsky, Elise G. Viox, Yiran Wang, Hui Wei, Yongping Yang, Amy F. Zhou, Rui Chen, Lu Yang, Diana G. Scorpio, Adrian B. McDermott, Lawrence Shapiro, Bridget Carragher, Clinton S. Potter, John R. Mascola, Peter D. Kwong

## Abstract

A central goal of HIV-1-vaccine research is the elicitation of antibodies capable of neutralizing diverse primary isolates of HIV-1. Here we show that focusing the immune response to exposed N-terminal residues of the fusion peptide, a critical component of the viral entry machinery and the epitope of antibodies elicited by HIV-1 infection, through immunization with fusion peptide-coupled carriers and prefusion-stabilized envelope trimers, induces cross-clade neutralizing responses. In mice, these immunogens elicited monoclonal antibodies capable of neutralizing up to 31% of a cross-clade panel of 208 HIV-1 strains. Crystal and cryo-electron microscopy structures of these antibodies revealed fusion peptide-conformational diversity as a molecular explanation for the cross-clade neutralization. Immunization of guinea pigs and rhesus macaques induced similarly broad fusion peptide-directed neutralizing responses suggesting translatability. The N terminus of the HIV-1-fusion peptide is thus a promising target of vaccine efforts aimed at eliciting broadly neutralizing antibodies.

---

Since crossing from chimpanzees ∼100 years ago^1^, HIV-1 has evolved to be one of the more diverse viruses to infect humans^2^. While antibodies capable of neutralizing ∼50% of circulating HIV-1 strains arise in half of those infected after several years^3^, the vaccine elicitation of antibodies capable of neutralizing divergent strains of HIV-1 remains an unsolved problem: antibodies elicited by current candidate vaccines fail to neutralize more than a small fraction of the diverse primary isolates that typify transmitted strains of HIV-1 (ref. ^4,5^).

We and others have isolated broadly neutralizing antibodies from HIV-1-infected donors and coupled antibody identification with structural characterization to delineate sites of vulnerability to neutralizing antibodies^6,7^. These antibodies target the viral entry machinery, the envelope (Env) trimer, composed of three gp120 and three gp41 subunits. Dozens of structurally defined epitopes have been determined that can be categorized into a handful of Env regions.

The majority of identified neutralizing antibodies have characteristics that may make them difficult to elict by vaccination, including those to the CD4-binding site^8,9^, where extensive somatic hypermutation (SHM) appears to be required^10–12^, those to a quaternary site at the trimer apex^13–16^, where unusual recombination appears to be required^13,14,17,18^, those to a glycan-V3 supersite^19–21^, where recognition of *N*-linked glycan appears to be required^20–22^, and those to the membrane-proximal external region^23–26^, where co-recognition of membrane^27–29^ and immune tolerance appear to be required^30^.

Recently, we identified an antibody, N123-VRC34.01 (ref. ^31^), named for donor (N123), lineage (VRC34) and clone number (01), and hereafter referenced without donor prefix. VRC34.01 targets primarily the conserved N-terminal region of the HIV-1 fusion peptide (FP), a critical component of viral entry machinery^32^. FP is composed of 15-20 hydrophobic amino acids at the N terminus of the gp41 subunit of HIV Env and had been thought to be poorly immunogenic: hidden from the immune system in the native prefusion state of Env and buried in the cell membrane after Env rearranges into the postfusion state. The VRC34.01 antibody, however, revealed the N-terminal half of FP to be a site of neutralization vulnerability. VRC34.01 directs the majority of its binding energy to eight N-terminal residues of FP, with the rest coming from interactions with Env including glycan N88 (ref. ^31^). The ability to neutralize HIV-1 through recognition of a linear peptide, which is both conserved in sequence and exposed in the prefusion-closed conformation of Env, suggested that the site of vulnerability defined by VRC34.01 might be amenable to epitope-based approaches to vaccine design. In this study we describe an antibody-to-vaccine development process. Beginning with the epitope of VRC34.01, we engineered immunogens with antigenic specificity for FP-directed antibodies, immunized C57BL/6 mice, and analyzed the resultant antibodies. Based on these analyses, we devised 2^nd^-generation immunization regimens that generated improved neutralizing monoclonal antibodies. We extracted insights from the murine immunizations and applied these to immunize guinea pigs and rhesus macaques. Overall, these vaccine studies demonstrated the ability, in multiple standard-vaccine test species, to induce serum responses capable of neutralizing a substantial fraction of primary isolate strains representative of the global diversity of HIV-1.

## Fusion peptide-based immunogens

The N-terminal eight residues of FP were chosen as a vaccine target, based on their recognition by antibody VRC34.01 in co-crystal structures and on molecular dynamics simulations, which showed these residues to be exposed and flexible in conformation^31^. To obtain immunogens capable of eliciting FP-directed antibodies, we utilized structure-based design to engineer FP-containing immunogens and assessed their antigenic specificity against a panel of antibodies encompassing both broadly neutralizing antibodies and poorly or non-neutralizing antibodies, with an emphasis on antibodies reported to engage FP as part of their recognized epitope such as ACS202 (ref. ^33^) and PGT151 (ref. ^34^) (Fig. 1a). We used an antigenicity score derived from the binding affinity of epitope-specific neutralizing and weak/non-neutralizing antibodies (see Methods) to estimate the epitope-specific suitability of each immunogen (Supplementary Fig. 1a). We produced epitope scaffolds that incorporated the N-terminal eight amino acids of FP, and in some cases, included added sites of *N*-linked glycosylation, which we positioned analogously to FP and glycan N88 in the VRC34.01 epitope. We characterized two scaffolds, which were trimeric and tetrameric from Protein Data Bank (PDB) 3HSH^35^, and 1SLF^36^, respectively, and also assessed epitope scaffolds engineered from PDB 1M6T^37^ and from PDB 1Y12 (ref. ^38^), which we previously described^31^ (Supplementary Fig. 1b-g), with the oligomeric scaffolds generally showing stronger binding to the FP-directed antibodies.

**Figure 1.**
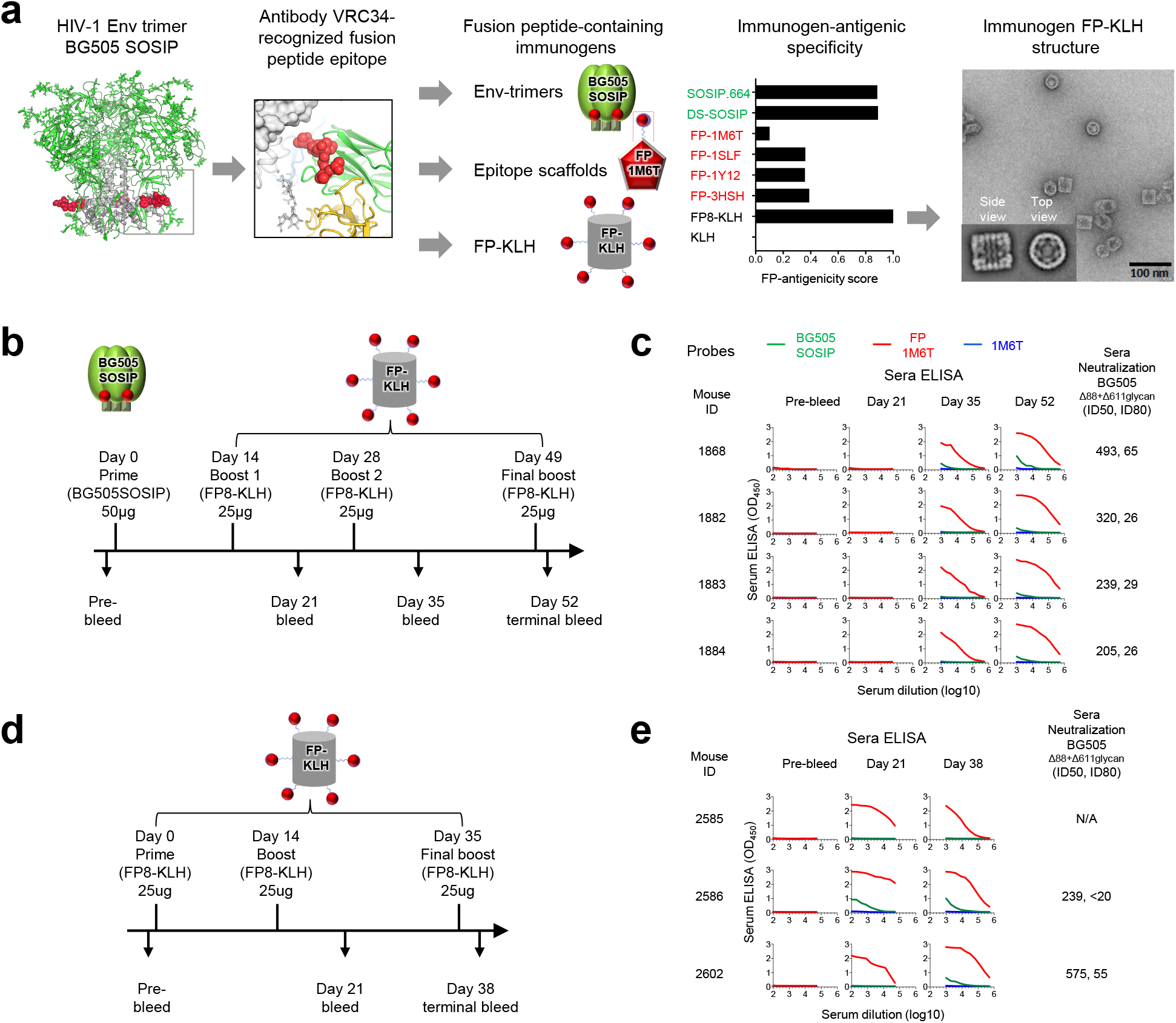
Design, properties, and immunogenicity of FP immunogens based on the epitope of antibody VRC34.01. (**a**) Structure-based design, antigenic characteristics, and EM structure of FP immunogens. The glycosylated structure of the HIV-1 Env trimer is shown at far left, with exposed N-terminus of FP highlighted in red. Subsequent images show site recognized by VRC34.01 antibody, schematics and antigenicity of FP immunogens, and negative stain EM of FP-KLH (see Supplementary Fig. 1 for details of FP antigenicity). For EM study, n=3 experiments were performed independent with similar results. (**b**) Immunization regimen 1. At day 52, mouse spleens were harvested and hybridomas created. (**c**) ELISA and neutralization of serum from regimen 1-immunized mice. Protein probes used for ELISA are defined in top row and include BG505 SOSIP.664 (green), FP-epitope scaffold based on PDB 1M6T (red), and 1M6T scaffold with no FP (blue). Column 1 defines mouse identification number and subsequent columns show ELISA and neutralization. ELISAs are shown as a function of serum dilution for pre-bleed, days 21, 35, and 52 (ELISA curves colored according to probe, with sera mostly unreactive with IM6T scaffold with no FP). Neutralization (ID_50_, ID_80_) values provided for day 52 serum; see Supplementary Fig. 2a for neutralization details. (**d**) Immunization regimen 2. At day 38, mouse spleens were harvested and hybridomas created. (**e**) ELISA and serum neutralization of serum from regimen 2-immunized mice, displayed as in **c**.

We also created a FP-carrier protein conjugate, by coupling the eight N-terminal residues of FP with an appended C-terminal cysteine to lysine residues in keyhole limpet hemocyanin (KLH), a carrier protein often used to improve immunogenicity^39^. The resultant FP8-KLH was robustly recognized by FP-directed antibodies and stable to extremes of temperature and osmolality, and at pH 10.0, but not at pH 3.5, and negative stain-EM revealed the FP-conjugated KLH to retain the barrel shape of KLH (Supplementary Fig. 1b, c, h, i). When assessed with the FP-antigenicity score (Fig. 1a), FP8-KLH was superior to the FP-epitope scaffolds and similar to the stabilized Env trimers, such as the SOSIP.664 (ref. ^40^) or DS-SOSIP^41^ trimers from the clade A strain BG505.

## Immunogens with high FP-antigenic specificity induce FP-directed neutralizing responses

To assess the ability of the 1^st^-generation epitope-based vaccine immunogens to elicit neutralizing responses, we tested immunization regimens using the two immunogens with the highest FP-antigenicity scores: FP8-KLH and stabilized Env trimer. In an initial experiment, four C57BL/6 mice each received 50 μg of BG505 SOSIP Env trimer and were boosted with 25 μg of FP8-KLH at day 14 (Fig. 1b). After a second boost at day 28, strong FP-ELISA responses were observed at day 35. After a final boost at day 49, we tested day 52 serum for neutralization of the Env-pseudovirus BG505, and also of a BG505 Env variants missing glycans at positions 88 or 611, as these Env variants are more sensitive to FP-directed antibodies^31^. While serum neutralization of wild-type BG505 generally did not pass our ID_50_ threshold for murine neutralization (at least 1:40 and at least 2-fold above background), we did observe unambiguous serum neutralization for the Δ88+611 glycan-deleted variant of BG505 by all four of the mice (Fig. 1c, Supplementary Fig. 2a).

In a subsequent experiment, three mice each received 25 μg of FP-KLH, followed by boosts at day 14 and day 35 (Fig. 1d). Serum ELISAs revealed Env trimer recognition in mouse 2586 at days 21 and 38, which also appeared in a second mouse after the third boost. We tested day 38 serum for neutralization and observed neutralization for the Δ88+611 glycan-deleted variant of BG505 by two of the sera (Fig. 1e, Supplementary Fig. 2a).

## First generation FP-directed antibodies neutralize up to ∼10% of HIV-1 strains

To provide insight into the antibodies elicited by FP-containing immunogens, we selected B cell hybridomas capable of binding both BG505 SOSIP trimer and FP-1M6T scaffold, using B cells from mouse 1868 (immunized with Env trimer and FP-KLH) and mouse 2586 (immunized with FP-KLH only). Sequences of eight hybridomas from mouse 1868 and five hybridomas from mouse 2586 revealed seven vaccine-elicited (v) antibody lineages (vFP1-vFP7), which could be segregated into three classes defined by similar B cell ontogeny and structural mode of recognition^6^, which we named vFP1, vFP5, and vFP6, after the first identified member of each class (Supplementary Fig. 3a, b, Supplementary Tables 1-2).

We tested neutralization of the 12 vFP antibodies on a panel of wild-type and glycan-deleted HIV-1 variants. Clear neutralization of wild-type HIV-1 strains was observed with only a few of the vFP1-class antibodies (vFP1.01, vFP7.04 and vFP7.05). We assessed the three best vFP1 antibodies along with antibody vFP5.01 on a well-characterized panel of 208 Env-pseudoviruses^5^, encompassing diverse strains of primary isolates from all of the major clades, of which 154 were resistant to neutralization by CD4-induced antibodies 17b and 48d, the V3-directed antibodies, 447-52D and 3074, and antibody F105, which recognized open conformations of Env (Supplementary Fig. 3c, Supplementary Table 3a). Notably, vFP1.01 neutralized 18 strains at 50 μg/ml (8.2% breadth on 208-strain panel; 7.7% breadth on 154-resistant strain panel), while vFP7.04 neutralized 20 strains at 50 μg/ml (9.6% breadth on 208-strain panel; 8.3% breadth on 154-resistant strain panel). vFP5.01, by contrast, neutralized only two strains, both sensitive to V3 and CD4-induced antibodies. Overall, neutralization by these 1^st^-generation monoclonal antibodies was of limited breadth and potency, although the best vFP antibodies could neutralize selected diverse strains of HIV-1.

## Disparate antibody-bound FP conformations

To provide insight into the structural basis for neutralization by the 1^st^-generation vaccine-elicited antibodies, we determined crystal structures for the antigen-binding fragment (Fab) of vFP1.01 and vFP5.01 antibodies in complex with the N-terminal eight residues of FP (Ala512-Phe519) at 2.0- and 1.5-Å resolution, respectively (Supplementary Table 4a, Supplementary Fig. 4, also see expanded views in Fig. 2a, b). The vFP1.01 co-crystals with FP were orthorhombic with four molecules per asymmetric unit, and in all four independent copies, Fab and FP assumed similar conformations, with FP adopting a curved structure, with no intrachain-backbone hydrogen bonds. The N terminus of FP (Ala512) was buried between heavy and light chains, with the amino terminus forming a buried salt bridge with Glu34vFP1.01-LC, which was germline encoded and shielded from solvent by a tetra-tyrosine cage, comprising tyrosines at residues 27D_vfp1.01-lc_, 32_vfp1.01-lc_, 96_vfp1.01-lc_, and 98_vfp1.01-hc_ (for clarity, we reference the molecule as a subscript for all molecules other than HIV Env by antibody name and HC or LC for heavy or light chain, respectively). The FP-main chain paralleled the curvature of the vFP1.01 CDR H3, albeit with opposite orientation, up to residue Ile515, which packed against the body of the heavy chain, before extending from antibody into the main body of the trimer with Gly516-Phe519 (Supplementary Fig. 4a, b).

**Figure 2.**
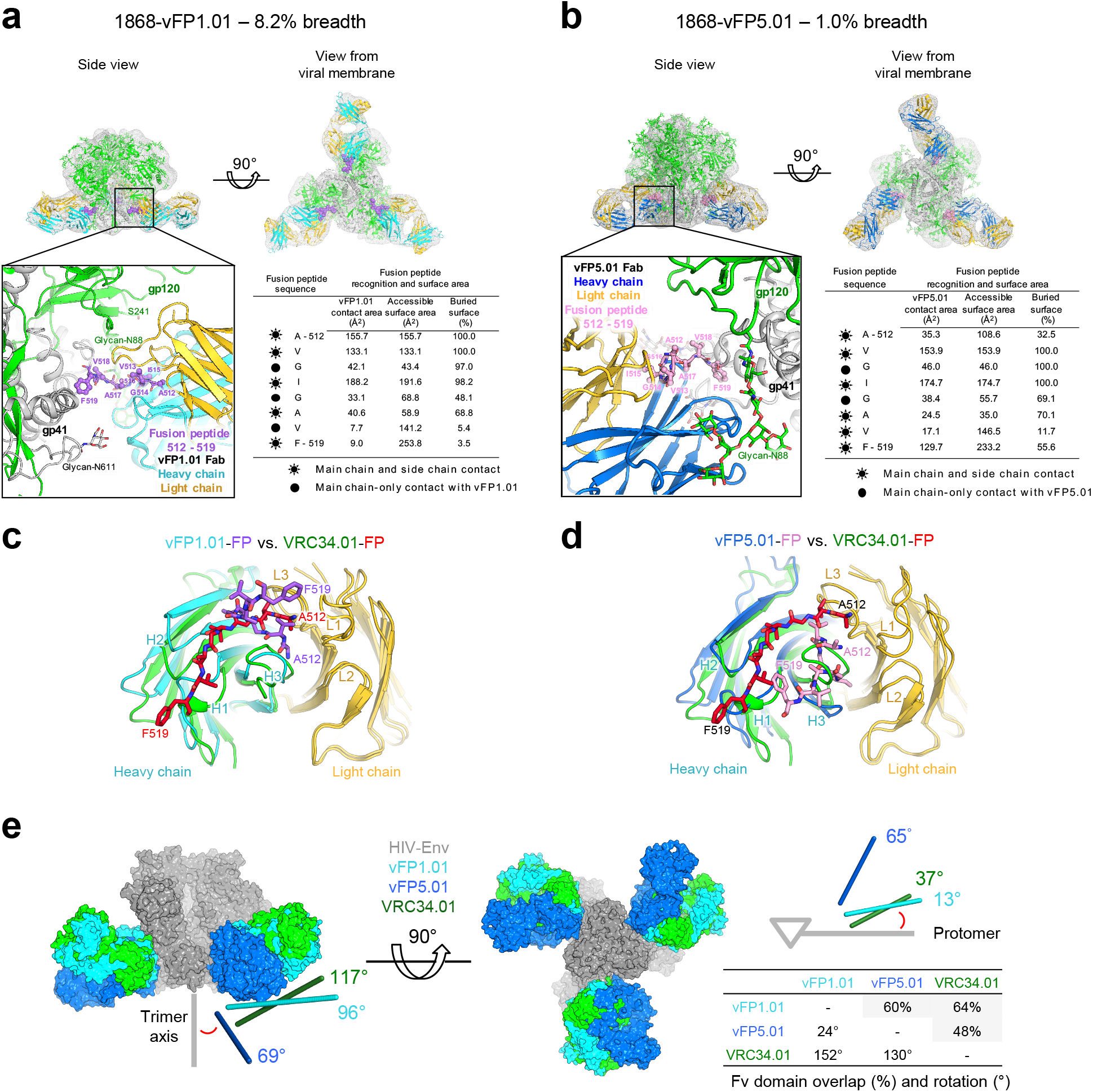
FP assumes disparate antibody-bound conformations, with neutralization restricted to a select angle of trimer approach. (**a**) Structural definition of vFP1.01 recognition. Top panels, cryo-EM reconstruction at 8.6 Å resolution (density shown in gray) of Fab vFP1.01 in complex with BG505 DS-SOSIP trimer. Expanded view, crystal structure of Env trimer and FP-bound Fab vFP1.01 at 2.0 Å resolution, as modeled into the cryo-EM map by rigid-body docking. Env trimer in green for gp120 and gray for gp41, Fab vFP1.01 in cyan for heavy chain and in yellow for light chain, and FP N-terminus in purple. Surface areas provided for N-terminal region of FP. (**b**) Same as a, but for vFP5.01 with FP in pink. Note that the angle of the lower right Fab differs from the angles of the other two. (**c**) Comparison of FP bound by vFP1.01 versus VRC34.01, with antibody shown in ribbons and FP in stick representation. (**d**) Same as c, but for vFP5.01. (**e**) Angle of recognition and Fv-domain overlap for vFP1.01, vFP5.01, and VRC34.01. Measured angles (red) are between antibody angle of approach and Env-trimer axis (left, with viral membrane located below trimer) and between antibody angle of approach parallel to viral membrane and Env protomer (right, looking down trimer axis towards viral membrane).

The vFP5.01 co-crystals with FP were monoclinic, with one molecule per asymmetric unit. vFP5.01 bound FP at the interface of heavy and light chains with the peptide adopting an overall hook structure: starting with a surface-exposed Ala512, dipping into the hydrophobic antibody interface with aliphatic side chains of Val513 and Ile515 anchoring the FP-N terminus, before turning at Gly516, and extending from antibody towards Env (Supplementary Fig. 4c, d).

Comparison of antibody-bound crystal structures indicated substantial differences in FP conformation (Fig. 2c, d). While antibodies VRC34.01, vFP1.01 and vFP5.01 recognized the N-terminus of FP by using a similar region at the CDR H3-CDR L3 interface of each antibody, the conformations of the antibody-bound FPs were substantially different: the vFP1.01-bound FP formed a U-shaped structure focused at the heavy-light interface, the vFP5.01-bound FP in an extended conformation to interact with CDR H3, and the VRC34.01-bound FP in an extended conformation to interact with CDR H1. To place these disparate antibody-recognized conformations of FP into a more general context, we used principal component analysis to cluster N-terminal FP conformations from a molecular dynamics simulation of fully glycosylated HIV-1 Env. Four prevalent clusters of fusion-peptide conformations were observed (Supplementary Fig. 4e-f), with FP-directed antibodies recognizing disparate but prevalent conformations of FP.

## Restricted angle of approach for FP-directed neutralization

To position the vFP1.01 and vFP5.01 structures with FP into the context of the HIV-1-Env trimer, we collected cryo-electron microscopy (cryo-EM) data for these FP-directed antibodies complexed to the BG505 SOSIP trimer. With Fab vFP1.01, approximately 14,000 particles yielded an 8.6-Å resolution reconstruction after three-fold averaging (Supplementary Fig. 5a-e); the resulting structure (Fig. 2a) showed three Fabs laterally interacting with the Env trimer. With Fab vFP5.01, several particle classes were observed yielding 14.7- and 19.6 -Å resolution reconstructions (Supplementary Fig. 5f-h); these asymmetric reconstructions indicated each of the vFP5.01 Fabs to approach Env differently (Fig. 2b).

To provide insight into recognition of the Env trimer, we analyzed the approach angle of the FP-recognizing antibodies (Fig. 2e). Overall, the approach of antibodies directed primarily to FP and capable of neutralizing diverse HIV-1 strains was similar, suggesting restrictions on trimer approach for effective FP-directed neutralization.

## Considerations for improved immunizations

Analysis of the 1^st^-generation antibodies indicated effective FP-directed neutralization to occur preferentially at a restricted angle of trimer approach, thereby suggesting that boosting with Env trimer might elicit improved neutralization. We sought additional clues from analysis of the 1^st^-generation vFP antibodies to improve FP immunization.

To provide insight into sequence requirements for neutralization, we created a panel of peptides comprising Ala and Gly mutants of the N terminus of FP and screened for recognition by vaccine-elicited antibodies and by VRC34.01 (Supplementary Fig. 6a, Supplementary Table 5). The Ala-Gly mutants only affected vFP1.01 recognition if they occurred within the first four residues of FP (512-515). For vFP5.01, a more extensive range was observed, with alterations to Ala-Gly at residues 513, 514, 515, 516, and 519 affecting recognition. VRC34.01 recognition by comparison was intermediate between vFP1.01 and vFP5.01, being sensitive to changes at 513, 515, and 516, and partially sensitive to changes at 518 and 519. These results indicated a preference for N-terminal residues for effective neutralization, thereby suggesting that N-terminal focusing might improve neutralization of the vaccine-elicited antibodies.

Although we did not observe a significant improvement in neutralization titers upon Env priming (Fig. 1), we nevertheless analyzed the degree of affinity maturation for vFP1-class antibodies as this would lend insight into the induction of these antibodies. We identified vFP1-class antibodies from three additional mice, two of them (1882 and 1883), primed with Env trimer, and one (2602), immunized with only FP-KLH (Supplementary Fig. 7a). Notably, vFP1-class antibodies primed with Env trimer showed higher somatic hypermutation (SHM) with statistically significance (Supplementary Fig. 7b). Thus, while Env-trimer priming did not improve neutralization, it did appear to prime vFP1-class antibodies.

## Second generation FP-directed antibodies neutralize up to 31% of HIV-1 strains

To elicit improved FP-directed antibodies, we tested 11 immunization regimens on 16 C57BL/6 mice (Supplementary Fig. 8a). The vaccine regimens included a BG505 SOSIP-trimer prime and various FP-KLH boosts, along with a BG505 DS-SOSIP boost on a subset of five of the mice. The FP-KLH boosts utilized different lengths of FP, ranging from FP6 through FP10, which incorporated 6 through 10 residues from the N-terminal FP sequence of strain BG505. Serum neutralization titers for the five trimer-boosted animals were especially improved, reaching an ID_50_ as high as 77,379 (mouse 2716) for the glycan deleted (Δ88+611) BG505, (Supplementary Fig. 8a,b). We tested the serum neutralization for the 5 trimer-boosted mice on a panel of 10 selected VRC34-sensitive wild-type strains, which encompassed divergent HIV-1 clades (Supplementary Figs. 2b and 8c). While modest neutralization from these five sera was observed against many of the selected strains, serum from mouse 2716 achieved ID_80_ neutralization against most of these viruses. Analysis by peptide competition indicated neutralization by the mouse 2716 serum to be targeted primarily to FP (Supplementary Fig. 2c).

For each of the 16 immunized C57BL/6 mice, we isolated and characterized monoclonal antibodies. These could be parsed into 21 lineages, vFP12-vFP32 (Supplementary Table 1a-c), and we screened each of the vFP antibodies against two wild-type viruses, the clade A BG505 and the clade B 3988.25; 24 antibodies showed wildtype neutralization, with most neutralizing 70% or more of the 10-strain panel (Supplementary Table 3b).

We chose two antibodies, 2712-vFP16.02 and 2716-vFP20.01, for in depth assessment (vaccine-elicited FP antibodies were named for mouse ID-lineage.clone, with antibody 2716-vFP20.01 being clone 01 from lineage vFP20 isolated from mouse ID 2716). Notably, on the 208-strain panel, these two antibodies achieved 31% and 27% neutralization breadth, respectively, when assessed at a maximum IC_50_ level of 50 μg/ml (Fig. 3, Supplementary Table 3a). The breadth and potency of these two vFP antibodies were similar to that of antibody 2G12 (Supplementary Fig. 11b), which has been shown to delay rebound and to induce selection pressure on the viral quasispecies when passively infused^42–44^. Neutralization extended to multiple clades, including clades A, B, and C, but for several clades, including clades AC, AG, D, CD, and G, no neutralization was observed. With the neutralized clades, the immunized FP8 sequence was generally present, whereas with the non-neutralized clades, this FP8 sequence was not present (with clade AE, the prevalent FP8 sequences included changes in the C-terminal region of FP8 (Supplementary Table 6), which were apparently tolerated by both vFP16.02 and vFP20.01 antibodies). We observed strong correlation between FP-directed antibody breadth on the 208-strain panel and the 58-strain panel with identical FP8 sequences, indicating overall breadth of the vFP antibodies to correlate with the ability to neutralize strains with the most prevalent FP8 sequence (Fig. 3, lower panels). These results provide proof-of-principle for the ability of FP-epitope focusing to induce FP-directed antibodies with promising neutralization breadth.

**Figure 3.**
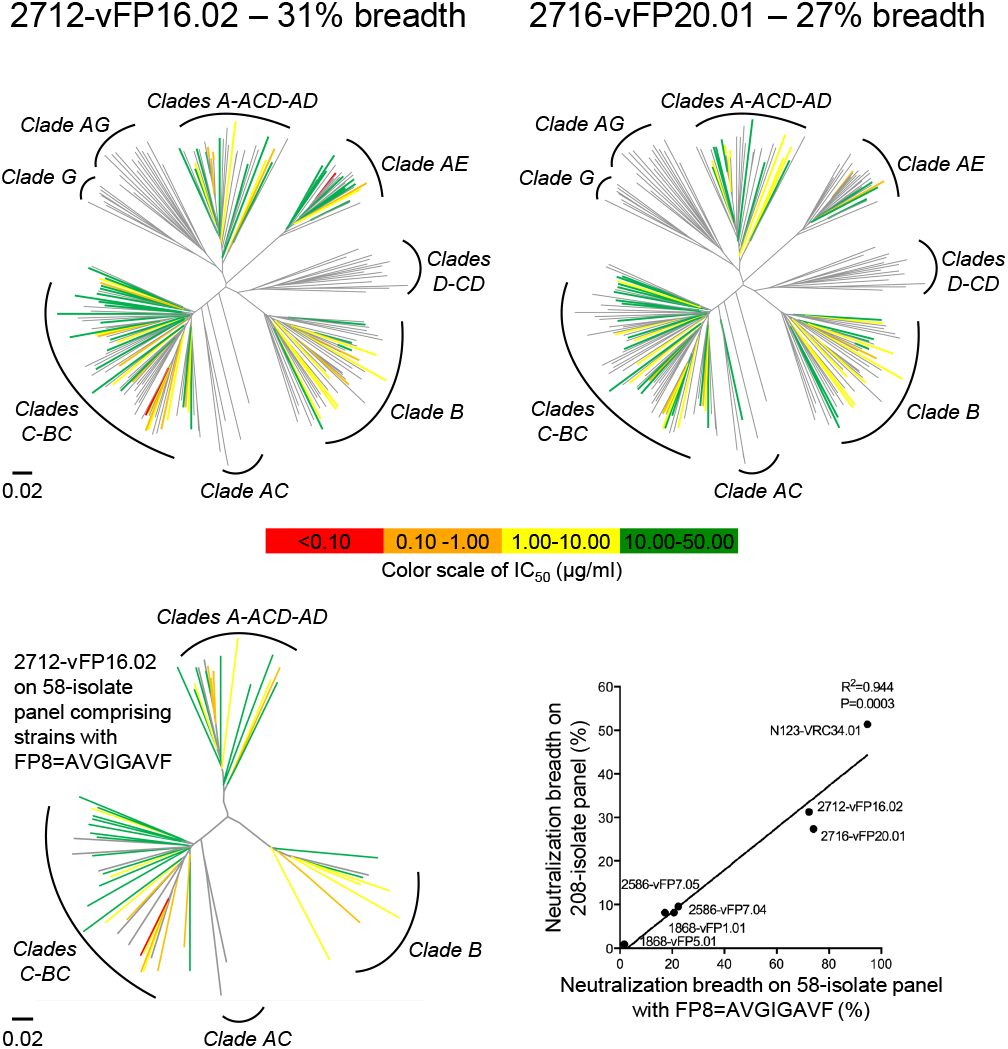
Second-generation vaccine-elicited antibodies neutralize up to 31% of HIV-1. Neutralization dendrograms display the diversity of tested viral strains, with branches colored according to neutralization potency (non-neutralized branches shown in gray). Top row: 208-strain panel for vaccine-elicited antibodies vFP16.02 and vFP20.01. Bottom left, 58-strain panel, displaying only branches with FP8=AVGIGAVF. Bottom right, comparison of breadth on 58- and 208-strain panels shown with Pearson correlation (n=7 antibodies).

## Structures and Env interactions of 2^nd^-generation antibodies, vFP16.02 and vFP20.01

To gain insight into the cross-clade breadth observed with vFP16.02 and vFP20.01 antibodies, we determined their crystal structures in complex with FP (Supplementary Fig. 4g, h) and their cryo-EM structures in complex with HIV-1 Env trimer (Fig. 4). The crystal structures with FP revealed highly similar recognition of residues 512-517, with the vFP1-class antibodies constraining little of the FP conformation beyond residue 517 (Supplementary Fig. 4g, h, Supplementary Table 4a). The cryo-EM data were collected on a quaternary complex with BG505 DS-SOSIP trimer bound by antibodies PGT122 and VRC03, in addition to vFP16.02 or vFP20.01; the added antibodies increased the size of the complex and provided fiducial markers allowing better particle visualization and alignment. The resultant reconstructions displayed resolutions of 3.6- and 3.8-Å, respectively, when calculated according to the FSC 0.143 gold-standard criterion with soft-edged masks from which flexible constant regions had been removed (Fig. 4, Supplementary Fig. 9a-f, Supplementary Table 4b). In the reconstructions, we observed electron density for the vFP16.02 and vFP20.01 antibodies focused around the fusion peptide, with density becoming weaker, farther from their fusion peptide site of recognition (Supplementary Fig. 10a-b).

**Figure 4.**
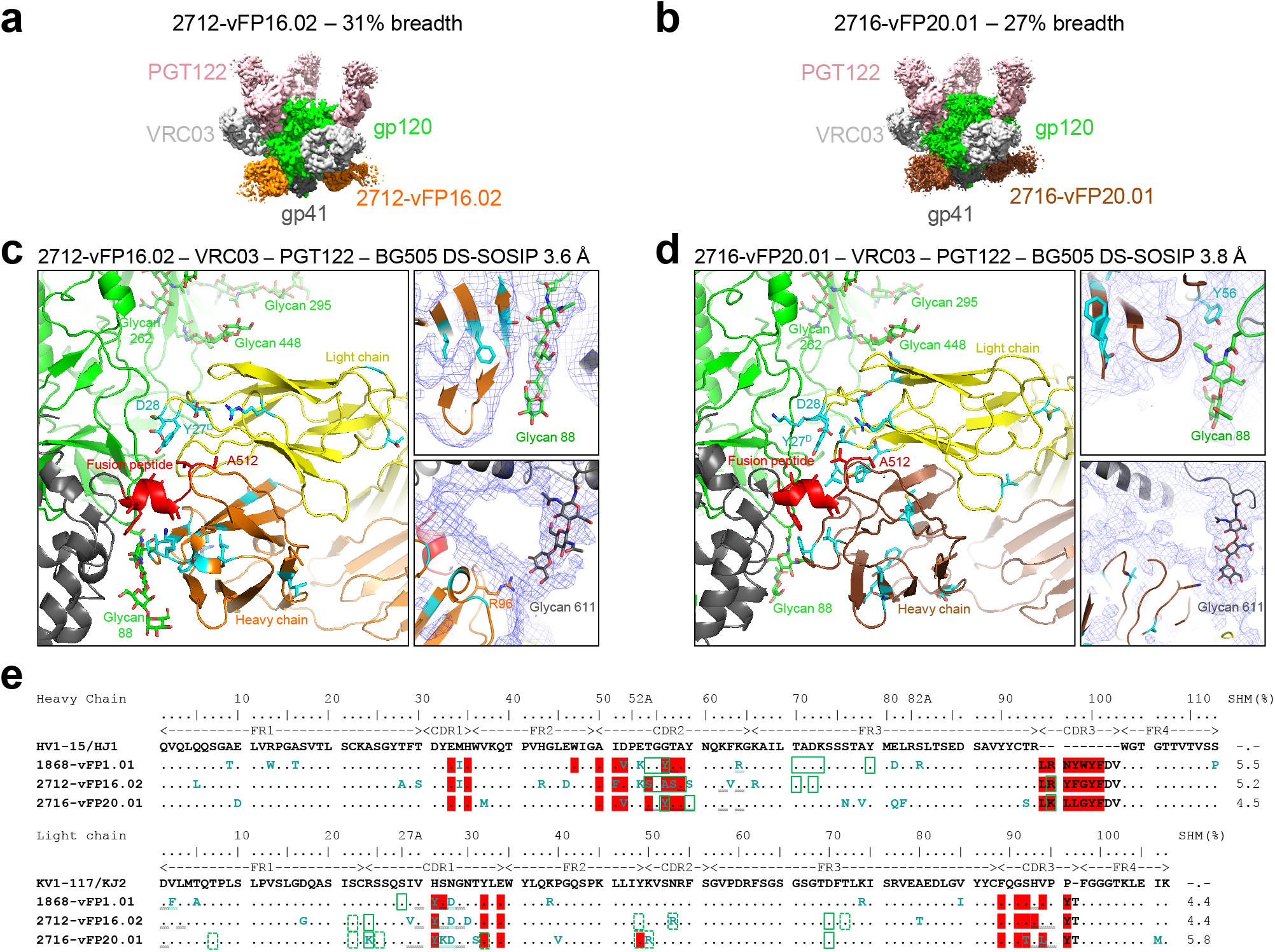
Substantial glycan contacts by 2^nd^-generation FP-directed antibodies. (**a**) CryoEM map of quaternary complex with antibody 2712-vFP16.02, segmented by components at a contour level that allowed visualization of Fv domains of antibodies (also see Supplementary Figs. 9 and 10). (**b**) Same as a, but with 2716-vFP20.01. (**c**) Details of vFP16.02 interaction, with right panels showing experimental EM density in blue mesh, with contour level adjusted to allow visualizing of partially ordered glycan. Residues altered by SHM highlighted in cyan. Antibody vFP16.02 contacts both FP and neighboring glycans to achieve 31% breadth. (**d**) Same as c, but for vFP20.01. (**e**) Sequence alignment of vaccine-elicited FP-directed antibodies and origin genes. FP contacts (red highlight), glycan contacts (green rectangle) and SHM (cyan font) are highlighted. Additional Env contacts are indicated by double underlining. Because the density from the cryo-EM reconstructions was not always sufficient to allow for atomic-level fitting, contacts shown with dotted green rectangles were inferred.

Glycan contacts between vFP antibodies and Env trimer were observed (Fig. 4c, d), and SHM was observed to occur preferentially at the interface with Env, especially involving interactions with FP and with *N*-linked glycan (Fig. 4e). In both antibody-Env complexes, glycans N448 and N611 displayed similar orientations, with glycan N448 buttressed by the light chain on one side and by glycan N295 on the other and with glycan N611 projecting from a neighboring Env protomer directly toward the antibody heavy chain. Glycan N88 also displayed ordered density for its protein-proximal sugars, though this differed in the two antibody complexes: in vFP16.02, substantial ordering was observed, with the glycan lodged between gp41 and the heavy chain (Fig. 4c); in vFP20.01, glycan N88 was less ordered and assumed a substantially different conformation to accommodate the SHM-altered Gly56Tyrvfp20.01-hc side chain (Fig. 4d). We note that even though removal of glycans neighboring FP (for example, glycan 611) improved neutralization by these antibodies, resistance analysis (Supplementary Table 7) indicated neutralization in the 208-strain panel to not be dependent on the absence of FP-neighboring glycans. Overall, the structures indicated murine vFP antibodies with promising breadth to substantially accommodate, if not partially recognize, FP-proximal *N*-linked glycan.

## Translation of murine vaccine regimens into guinea pigs and NHPs

We extracted insights from the murine immunization regimens, such as trimer boost and FP N-terminal focusing, and applied these to guinea pigs and rhesus macaques (non-human primates; NHPs). We immunized five guinea pigs using a scheme modeled on the mouse 2716 immunization regimen (Fig. 5a). At week 28, we observed 4 of 5 guinea pigs to show some heterologous virus neutralization on the 10-strain panel. When compared to DS-SOSIP alone immunized guinea pigs (Supplementary Fig. 7e), the 10-strain breadth induced by FP-KLH prime DS-SOSIP boost was significantly higher than induced by DS-SOSIP alone (Fig. 5a, b).

**Figure 5.**
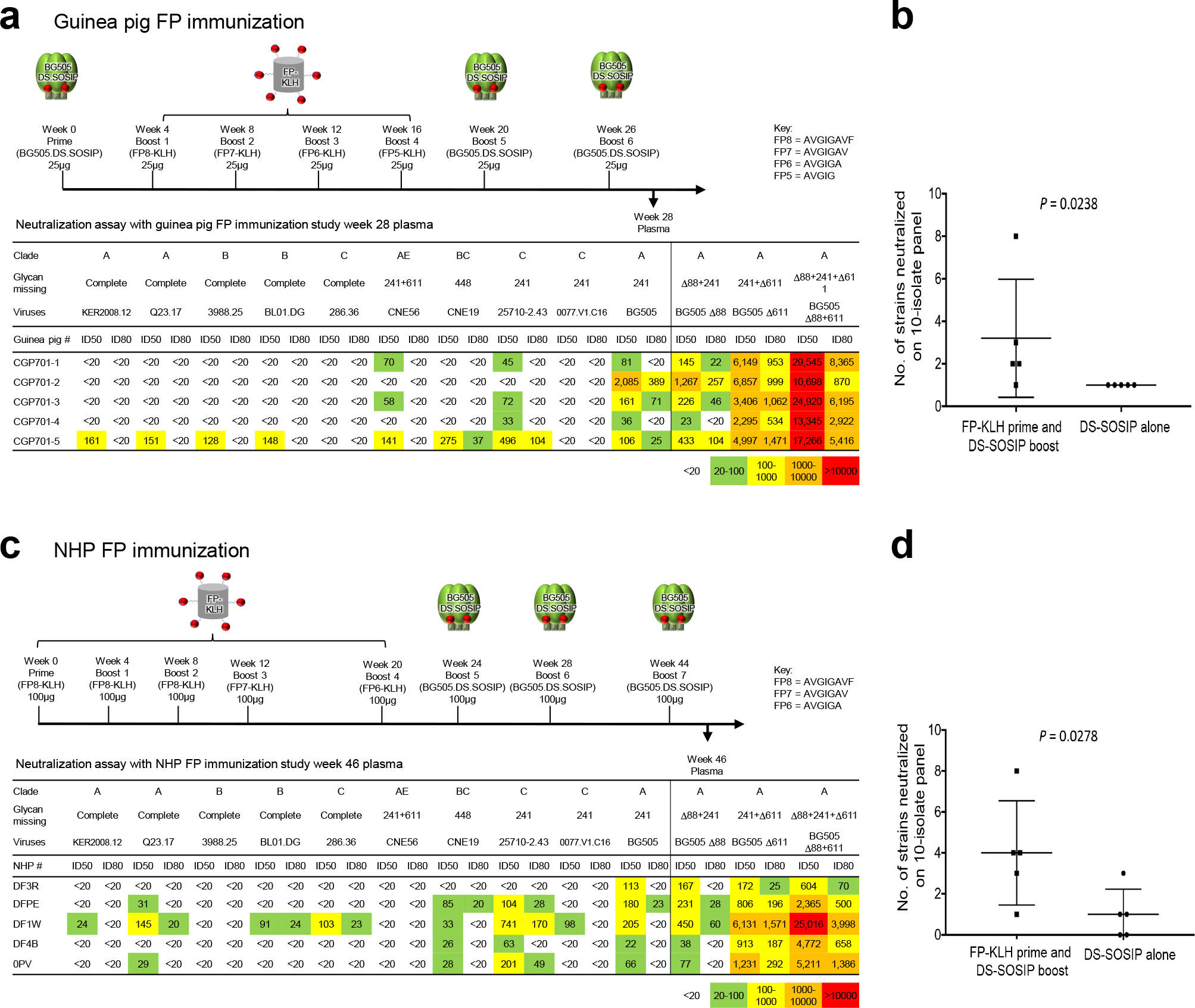
Immunization of guinea pigs and NHPs with FP-coupled carriers and DS-SOSIP trimer elicits heterologous neutralizing responses. (**a**) Elicitation of serum neutralizing responses in guinea pigs. Immunization regimen and week 28 serum ID_50_ titers as measured on a 10-wildtype strain panel, 5 with complete glycans around FP, and 5 naturally missing glycans at sites defined in the figure. Also shown are titers for Δ88, Δ611, and Δ88+611 glycan-deleted variants of BG505. (**b**) Plot comparing guinea pig-serum neutralization breadth for FP-KLH prime and DS-SOSIP-trimer boost regimen versus DS-SOSIP alone regimen (one-tailed Mann-Whitney; see Supplementary Fig. 7 for DS-SOSIP alone immunizations at 0, 4 and 16 weeks); n=5 animals for each group. (**c**) Elicitation of heterologous neutralizing responses in rhesus macaques. Immunization scheme, and week 46 serum titers assessed and displayed as in **a**. (**d**) Plot comparing NHP-serum neutralization breadth for immunization regimens, displayed as described in **b** for guinea pigs; n=5 animals for each group.

For NHP, we also used a regimen similar to that used for mouse 2716, but omitting the initial trimer prime and using three trimer boosts, so that we could compare to NHPs immunized with trimer only. We used a total of 5 FP-KLH primes with FP peptides of lengths 8-8-8-7-6. At week 46, we observed 4 of 5 NHP plasma to show heterologus virus neutralization on the 10-strain panel (Fig. 5c, d). As seen in the GP immunization study, this breadth of neutralization by NHP plasma was higher than that induced by immunization with DS-SOSIP alone (Supplementary Fig. 7f)^45^.

We further assessed all 5 NHP week 46 plasma on a 58-strain subset of the 208-strain panel, restricted to strains for which the sequence of FP was AVGIGAVF, the most prevalent FP sequence and the sequence in the FP8-KLH and BG505 DS-SOSIP immunogens. At a 1:20 dilution, 3 of 5 NHP plasma showed substantial neutralization breadth, with the broadest, NHP DF1W, achieving ID_50_ neutralization on 41 of 58 strains (70%) and ID_80_ neutralization of 13 of 58 strains (22%) (Fig. 6a, Supplementary Table 3c). Neutralization competition of the NHP DF1W plasma tested against neutralization-resistant HIV-1 strains with complete glycan around FP from four different clades indicated that a linear FP, matching the immunogen sequence, could adsorb virtually all of the neutralizing activity (Fig. 6b, Supplementary Fig. 2d). A dendrogram of these 58-Env strains (Fig. 6c) indicated neutralization to be distributed over several clades, consistent with the distribution of the immunized FP8 sequence in HIV-1. Additionally, the neutralization fingerprints of the three NHP plasma with substantial breadth clustered next to the neutralization fingerprints of the murine vFP1-class antibodies (Fig. 6d), providing a strong indication for the similarity of induced murine and NHP immune responses.

**Figure 6.**
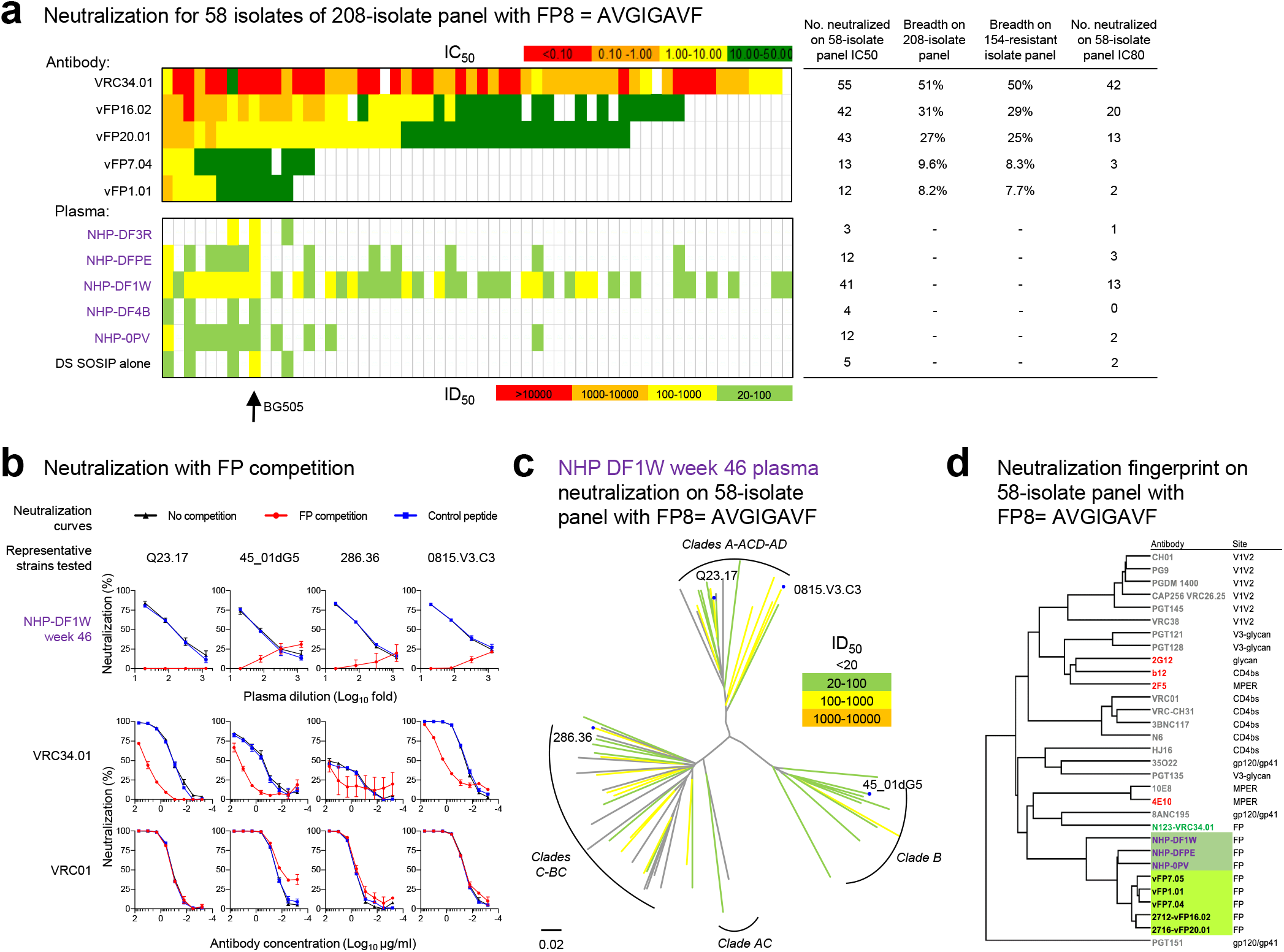
Patterns of neutralization indicate FP-directed responses in mice and NHP are related. (**a**) Neutralization on a 58-strain panel, comprising strains from the 208-strain panel with the FP sequence, AVGIGAVF, matching both FP8-KLH and BG505 trimer immunogens. Top panel, IC_50_ values for FP-directed antibodies. Bottom panel ID_50_ titers for NHP plasma: top rows, FP-KLH prime and DS-SOSIP-trimer boost regimen at week 46, 2 weeks after third trimer boost; bottom row, DS-SOSIP alone regimen at week 18, 2 weeks after third trimer boost. Number of neutralized strains and neutralization breadths are shown. (IC and ID provided in Supplementary Table 3c and Supplementary Fig. 7.) (**b**) Neutralization curves of NHP plasma (DF1W week 46) and control monoclonal antibodies (VRC34.01 and VRC01) on four representative strains in the presence of no peptide (black), FP (red) or an irrelevant Flag peptide (blue). Mean and standard deviation of results from triplicated experiments shown (n=3 independent experiments). Location of representative tested strains are labeled and shown on dendrogram in c. (**c**) Neutralization dendrogram (ID_50_) for NHP DF1W week 46 plasma on 58-strain panel. (**d**) Neutralization-fingerprint dendrogram calculated from 58-strain panel. Vaccine-elicited vFP antibodies (highlight with green background) and three NHP week 46 plasma (highlighted with forest green background for three of five NHP with sufficient neutralization to yield accurate fingerprint analysis) clustered next to each other (see also fingerprint dendrogram on 132-curated strains shown in Supplementary Fig. 11a).

## Discussion

The vaccine elicitation of antibodies capable of neutralizing diverse strains of HIV-1 has been a goal of HIV-1 research for over 30 years. While substantial strides have been made in the creation of prefusion-stabilized Env trimers^40,41,46–49^, responses elicited by these trimers have been primarily strain-specific. Here, we show that focusing the immune response to the exposed N-terminal residues of FP elicits antibodies of promising neutralization breadth in multiple vaccine-test species. Several factors contributed to the successful elicitation of neutralizing antibodies. First, antibody VRC34.01 (ref. ^31^) defined a precise target – not the entire FP, but only exposed N-terminal residues – upon which to focus. Second, glycan-deleted viruses^50^ provided a readout sensitive enough to detect initial weak serum responses. Third, we found prefusion-closed Env trimers^40^ to be necessary for boosting serum responses. Most important, however, may be the characteristics of the target site – the FP N-terminus – a conserved and exposed site of vulnerability, for which high conformational variability was compatible with broad neutralization (Supplementary Figs. 4e and 11c-g).

FP-directed antibodies of substantial breadth have been reported for other viral pathogens, including influenza A virus^51–52^, Ebola virus^53–55^ and Lassa virus^56^. Although the fusion peptides of these pathogens are substantially different from that of HIV-1, the generality of FP-targeted neutralizing antibodies suggests that characteristics of FP may be especially suited to vaccine targeting. Such characteristics include its high conservation (likely related to function), its chemistry (hydrophobic residues afford high binding energy), and its exposure (FP is often proximal to a site of proteolytic activation).

Is there a neutralization limit or constraint for vFP antibodies based on viral properties (e.g. related to strain-specific conformation or accessibility) other than the diversity of FP itself? Strong correlation was observed between Env trimer affinity and neutralization breadth (Supplementary Figs. 6b, c, 11d), suggesting that enhanced Env-trimer affinity would lead to increased breadth. Indeed, at 10-fold higher concentration (500 μg/ml), vFP16.02 neutralized 97% (56 of 58 strains) with the immunized FP8 sequence, and 47% of the 208-strain panel (97 of 208 strains) (Supplementary Table 3a). Thus, for strains with the same sequence as used in the FP immunizations, there did not appear to be an intrinsic limit to elicited FP-directed breadth.

Finally, is the observed elicitation of vFP-neutralizing antibodies in animal models translatable to humans? While broadly neutralizing antibodies against HIV-1 have been induced in animals such as cows^57^ and llamas^58^, these antibodies rely on species-specific characteristics^59,60^ (knob domains for cow and unpaired heavy chains for llama), which are not present in human antibodies. By contrast, there does not appear to be an intrinsic barrier to eliciting vFP antibodies in humans: genetic analysis indicates humans to have V-genes with some similarity to the germline genes of the murine vFP1-antibody class (Supplementary Fig. 12a, b); FP-directed antibodies can often be detected in HIV-1-infected donors by ELISA^31^; and FP-directed antibodies showed almost no indication of polyreactivity (Supplementary Fig. 12c, d). Overall, our results provide proof-of-principle for the ability of epitope-based vaccine design to induce FP-directed antibodies with neutralization breadth and indicate the exposed N terminus of FP to be a site of exceptional HIV-1 vaccine promise.

## ACKNOWLEDGMENTS

We thank J. Chrzas, U. Chinte, Z. Jin and staff at SER-CAT (Southeast Regional Collaborative Access Team) for help with x-ray diffraction data collection, B. DeKosky for suggestions on lineage discrimination, W. Rice and SEMC staff for assistance with cryo-EM data collection, and J. Stuckey for assistance with figures. We thank D. Burton and M. Fineberg for antibodies from International AIDS Vaccine Initiative’s (IAVI’s) Neutralizing Antibody Consortium (NAC) including PGT122 used in cryo-EM studies, B. Haynes for information on antibody CH07, R. Sanders for information on antibody ACS202, B. Graham for murine antibody 5C4, and the WCMC/AMC/TSRI HIVRAD team for their contributions to the design and validation of near-native mimicry for soluble BG505 SOSIP.664 trimers. We thank members of the Structural Biology Section, Structural Bioinformatics Core Section, and Human Immunology Section of the Vaccine Research Center for helpful comments. Support for this work was provided by the Intramural Research Program of the Vaccine Research Center, National Institute of Allergy and Infectious Diseases, National Institutes of Health. This work was also supported in part IAVI’s NAC (J.R.M and P.D.K.) and with federal funds from the Frederick National Laboratory for Cancer Research, NIH, under contract HHSN261200800001E (Y.T.). I.S.G. received support from NIH grant R01 AI131722. B.C. and C.S.P. received support from NIH grant GM103310 and from the Simons Foundation (349247). Use of insertion device 22 (SER-CAT) at the Advanced Photon Source was supported by the U.S. Department of Energy, Basic Energy Sciences, Office of Science, under contract W-31-109-Eng-38.

## AUTHOR CONTRIBUTIONS

K.X. conceived and led the project, as well as determined all crystal structures; P.A. determined cryo-EM structures; R.K. and N.A.D.-R. coordinated neutralization assessments; C.C. coordinated guinea pig and NHP immunization; G.-Y.C. coordinated statistical and bioinformatical analysis; K.L. prepared proteins and co-determined crystal structures; M.K.L. and R.T.B. assessed antibody neutralization in the large panel; C.-H.S. performed antibody sequence analysis; M.S. performed all Alanine/Glycine scans; B.Z. prepared antibodies for large panel, and performed various binding analysis; T.Z. performed SPR analysis, calculated antibody approaching angel to trimer and made antibody neutralization dendrograms; M.A. performed antibody autoreactivity test; I.S.G. performed antibody neutralization finger-print analysis; T.L. performed molecular dynamics analysis; S.D., K.M., C.W.C., E.G.V. and A.P.R. co-performed neutralization assay; A.D., D.P., B.Z. and Y.Y. helped with protein expression; E.T.E., V.P.D. and H.W. helped with cryo-EM structures; X.C., H.G., J.G., M.S., and Y.D.K. performed protein purification; K.R.H, A.J.J., K.E.F, D.G.S, J.-P.T assisted in NHP study; Y.-T.L. and Y.W. assisted with X-ray crystal datasets processing; B.Z., L.O. and M.C. helped with immunogen preparation and characterization; R.R. and K.F. performed statistical and bioinformatical analysis; Z.S. performed antibody gene comparison; Y.T. performed negative-stain EM; T.Y.O participated in immunogen binding test; A.F.Z. helped with SPR assay; R.C. and L.Y. supervised the research team in GenScript; A.B.M supervised MSD-based immunogen antigenicity characterization; L.S. supervised antibody gene analysis and comparison; B.C. and C.S.P. supervised cryo-EM studies; K.X., L.S., J.R.M. and P.D.K. wrote the manuscript, and all authors read, edited and approved the manuscript. J.R.M. and P.D.K. supervised the study.

## METHODS

### Peptide synthesis and peptide-carrier protein conjugate preparation

HIV-1 fusion peptides (FPs) were synthesized (GenScript) with a free amine group on the N terminus. FP His-tagged peptides were synthesized (GenScript) with a six histidine residues tag at the C terminus of FP. To prepare peptide-carrier protein conjugates, each peptide with a cysteine residue added to the C terminus was conjugated to the carrier protein keyhole limpet hemocyanin (KLH) (Thermo-Scientific) using m-maleimidobenzoyl-N-hydroxysuccinimide ester (MBS) following the manufacturer’s protocol.

### Protein expression and purification

BG505 SOSIP, BG505 DS-SOSIP and their glycan-deficient variants were expressed and purified as described previously^41^. FP-epitope scaffold proteins, including FP-1M6T, FP-1Y12, FP-3HSH and FP-1SLF were expressed and purified as described previously^31^. vFP1.01, vFP7.04, vFP16.02 and vFP20.01 antibodies used for structure determination were prepared as below. Heavy chain plasmids, encoding the chimera of mouse variable domain and human constant domain, with HRV3C cleavage site in the hinge region; and light chain plasmids, encoding the chimera of mouse variable domain and human constant domain were cotransfected in Expi293F cells (Thermo Fisher) using Turbo293 transfection reagent (SPEED BioSystem) according to the manufacturer’s protocol. Transfected cells were incubated in shaker incubators at 120 rpm, 37°C, 9% CO_2_ overnight. On the second day, one tenth culture volume of AbBooster medium (ABI scientific) was added to each flask of transfected cells and cell cultures were incubated at 120 rpm, 33°C, 9% CO_2_ for an additional 5 days. 6 days post-transfection, cell culture supernatants were harvested. IgGs were purified from the supernatants using protein A chromatography: after PBS wash and low pH glycine buffer elution, the eluate was immediately neutralized using 10% volume of 1M Tris buffer pH 8.0. Fabs were obtained either by HRV3C cleavage or papain digestion. The fragmented Fabs were further purified by size exclusion chromatograph (SEC) in a Superdex 200 column (GE) with a buffer containing 5mM HEPES, pH 7.5, 150mM NaCl.

### Negative-stain electron microscopy

Samples were diluted with a buffer containing 20 mM HEPES, pH 7.0, 150 mM NaCl, adsorbed to a freshly glow-discharged carbon-film grid, washed with the above buffer, and stained with 0.7% uranyl formate. Images were collected semi-automatically at a magnification of 100,000 using SerialEM^61^ on a FEI Tecnai T20 microscope equipped with a 2k x 2k Eagle CCD camera and operated at 200 kV. The pixel size was 0.22 nm/px. Particles were picked manually using the swarm mode in e2boxer from the EMAN2 software package^62^. Reference-free 2D classification was performed using EMAN2 and SPIDER^63^.

### Antigenic characteristics of fusion peptide immunogens with Biolayer Interferometry and MSD-ECLIA

Antigenic characteristics of KLH-coupled fusion peptide, FP scaffolds and BG505 Env trimers to various FP-targeting and Non-FP-targeting HIV antibodies were assessed with Biolayer Interferometry on an Octet RED384 (ForteBio) instrument and with MSD-ECLIA as previously described^41^.

### Antigenicity score

Antigenicity scores for the FP immunogens were calculated according to the metric defined in Supplementary Fig. 1a. This metric sums the average binding of neutralizing antibodies versus poorly/nonneutralizing antibodies, with binding defined as a function of the logarithm of antigen-binding affinity relative to upper and lower binding limits. This summation is weighted by the site targeted. An antigenicity score of 1for an FP immunogen therefore indicates both high specificity and tight affinity for FP-directed neutralizing antibodies; An antigenicity score of 0 for an FP immunogen would indicate either low specificity or weak affinity for FP-directed neutralizing antibodies. In Fig. 1a, the antigenicity score was calculated with only FP-directed antibodies; we would anticipate the FP-antigenicity score for trimer immunogens to decease relative to FP-specific immunogens such as FP8-KLH if non-FP-directed antibodies were considered.

### Mouse immunization (GenScript)

Female mice (C57BL/6) around 8 weeks old were immunized in two-week intervals with either HIV-1 Env trimer or FP-KLH, using either Adjuplex as adjuvant (Sigma-Aldrich Inc, MO) for trimer or GS-adjuvant (GenScript) for FP-KLH. 50 μg of immunogens were used for prime immunization and 25 μg immunogens were used in boost immunization. Intraperitoneal (IP) route was used for all mice immunizations. Sera samples were collected either 7 days or 14 days after each immunization for ELISA and other analyses.

All experiments were performed in accordance with protocols reviewed and approved by the Genscript’s Institutional Animal Care and Use Committee (IACUC, #ANT14-003 and #ANT17-003). All mice were housed and cared for in a facility in GenScript accredited by Association for Assessment and Accreditation of Laboratory Animal Care International (AAALAC International).

### Hybridoma creation and monoclonal antibody production

Terminal boosts were performed on the top responders from each immunization scheme as assessed with ELISA against the immunogens, three weeks after the last immunization. Mice spleens were harvested three days post terminal boost, and hybridomas were generated for monoclonal antibody selection following the standard procedure at GenScript. Monoclonal antibody selection was based on affinity to FP-scaffold (FP-1M6T) and BG505 Env trimer as measured by ELISA.

### Guinea pig and NHP protocols and immunizations

For immunization studies, all animals were housed and cared for in accordance with local, state, federal, and institute policies in an American Association for Accreditation of Laboratory Animal Care-accredited facility at the Vaccine Research Center, NIAID, NIH or at a contract facility (Bioqual Inc, MD). All animal experiments were reviewed and approved by the Animal Care and Use Committee of the Vaccine Research Center, NIAID, NIH, and covered under protocol VRC-13-431, VRC-16-667.

Female Hartley guinea pigs with body weights of 300 grams were purchased from Charles River Laboratories, MA. For each immunization, 400 μl of immunogen mix, containing 25 μg of specified, filter-sterilized protein immunogen and 80 μl of Adjuplex (Sigma-Aldrich Inc, MO or Adjuplex equvalent formulated at VRC) in PBS, was injected into the muscle of the two hind legs. While animals were under anesthesia, blood was collected through retro-orbital bleeding for serological analyses.

Female and male Indian origin rhesus macaques with body weights of 2-9 kg were used for immunization studies. For each immunization, 1 ml of immunogen mix, containing 100 μg of specified, filter-sterilized protein immunogen and 200 μl of Adjuplex (Sigma-Aldrich Inc, MO or Adjuplex equvalent formulated at VRC) in PBS, was injected via a needle syringe into the caudal thighs of the two hind legs. Blood was collected two weeks post immunization for serological analyses.

### ELISA

Fusion peptide ELISAs: Costar^®^ High Binding Half-Area 96-well plates (Corning, Kennebunk, ME) were coated with 50 μl/well of 2 μg/ml scaffold proteins in PBS overnight at 4°C. Between steps, plates were washed 5 times with PBS-T (PBS + 0.05% Tween) and incubated at 37°C for 1 hour. After coating, plates were blocked with 100 μl/well of blocking buffer (B3T: 150 mM NaCl, 50 mM Tris-HCl, 1 mM EDTA, 3.3% fetal bovine serum, 2% bovine albumin, 0.07% Tween 20, 0.02% Thimerosal). 2-fold serial dilution of 1:100 pre-immunization and 1:1000 post immunization sera were used in the ELISA. Goat anti-mouse IgG (HRP-conjugated, GenScript) at 1:5000 dilution was used for detection. Plates were developed with tetramethylbenzidine (TMB) substrate (SureBlueTM, KPL, Gaithersburg, MD) for 10 minutes before adding 1 N sulfuric acid (Fisher Chemical) to stop the reaction. Plates were read at 450 nm (Molecular Devices, SpectraMax using SoftMax Pro 5 software) and the optical densities (OD) were recorded.

BG505 SOSIP D7324 Capture ELISAs: Costar^®^ High Binding Half-Area, 96-well plates (Corning, Kennebunk, ME) were coated with 50 μl/well of 2 μg/ml of sheep D7324 antibody (AALTO Bio Reagents) in PBS overnight at 4°C. Between steps, except for addition of trimer, plates were washed 5 times with PBS-T (PBS + 0.05% Tween) and incubated at room temperature (RT) for 1 hr. After coating, plates were blocked with 100 μl/well of blocking buffer (5% Skim Milk, 2% bovine albumin, 0.1% Tween 20 in TBS). Next, 50 μl/well of 0.5 μg/ml D7324-tagged BG505.SOSIP trimer diluted in 10% fetal bovine serum in PBS was added and incubated at RT for 2 hours2-fold serial dilution of 1:100 pre-immunization and 1:1000 post immunization sera were used in the ELISA. Goat anti-mouse IgG (HRP-conjugated, GenScript) at 1:5000 dilution was used for detection. Plates were developed with tetramethylbenzidine (TMB) substrate (SureBlueTM, KPL, Gaithersburg, MD) for 10 minutes before adding 1 N sulfuric acid (Fisher Chemical) to stop the reaction. Plates were read at 450 nm (Molecular Devices, SpectraMax using SoftMax Pro 5 software) and the optical densities (OD) were recorded.

ELISA responses were plotted using PRISM (PRISM 7 GraphPad Software for Mac OS X).

### Genetic assignment of antibodies

Antibody sequences were submitted to the ImMunoGeneTics information system^®^ (IMGT, http://www.imgt.org) and subjected to variable(V), diverse(D) and joining(J) genes identification by alignment with the mouse germline sequences of the IMGT reference directory, and IMGT/JunctionAnalysis for a detailed analysis of the V-J and V-D-J junctions. We only considered confirmed functional germline genes for the assigned germline. Clustal Omega software was used to prepare multiple sequence alignment of antibody sequences for maximum likelihood phylogenetic tree construction using DNAML program in the PHYLIP package version 3.69 (http://evolution.genetics.washington.edu/phylip.html). Calculations were performed based on empirical base frequencies with transition/transversion (Ti/Tv) ratio of 2.0. Dendroscope 3 (dendroscope.org) was used to visualize phylogenetic trees^64^. The amino acid sequence alignments were visualized using BioEdit v7.2.5 editing software^65^. To calculate the minimal mutations required to switch between two different unmutated common ancestors, the unmutated common ancestor sequence was prepared by reverting the assigned V(D)J gene sequences into their corresponding germline sequences. Differences between unmutated common ancestor sequences were counted as the minimal mutations required to switch from one unmutated common ancestor to another.

### Antibody alanine/glycine scan

Binding of the vaccine elicited mouse vFP antibodies to sixteen His-tagged fusion peptide (residue 512-521), including wildtype and alanine/glycine mutants, was assessed using a fortéBio Octet Red384 instrument. Briefly, the sixteen peptides at 50 μg/ml in PBS were loaded onto Ni-NTA biosensors using their C-terminal histidine tags for 60 s. Typical capture levels were between 1.1 and 1.3 nm and variability within a row of eight tips did not exceed 0.1 nm. These peptide-bound biosensors were equilibrated in PBS for 60 s followed by capture of the antigen binding fragments (Fabs, 250 nM) of the vaccine elicited vFP antibodies, VRC34.01 and an RSV F antibody Motavizumab for 120 s and a subsequent dissociation step in PBS.

In all Octet measurements, parallel correction to subtract systematic baseline drift was carried out by subtracting the measurements recorded for a loaded sensor incubated in PBS. Data analysis was carried out using Octet software, version 9.0. The normalized responses obtained from one or triplicate dataset were plotted using PRISM (PRISM 7 GraphPad Software for Mac OS X).

### Surface plasmon resonance assay

Binding affinities and kinetics of antibodies to HIV-1 DS-SOSIP trimers and His-tagged fusion peptide were assessed by surface plasmon resonance on a Biacore T-200 (GE Healthcare) at 25 °C. To test antibody binding with HIV-1 DS-SOSIP trimers, 2G12 IgG was first immobilized on flow cells of a CM5 chip at ∼3000-8000 response unit. BG505 DS-SOSIP trimer and its glycan-deleted mutants, BG505 DS-SOSIP.Δ88 and BG505 DS-SOSIP.Δ611, at 500 nM in HBS-EP+ buffer (10 mM HEPES, pH 7.4, 150 mM NaCl, 3 mM EDTA and 0.05% surfactant P-20) were then captured onto 2G12 of one flow cell by flowing the protein solution for 60 s at a flow rate of 6 μl/min. Binding affinities for the 2G12-captured trimer were determined by using a serial dilution of antibody Fab solutions starting at 200nM during association phase. A dissociation phase at 30 μl/min for 300 s was used to determine binding kinetics. The surface was regenerated by flowing 3M MgCl_2_ solution for 30 s at a flow rate of 50 μl/min. Blank sensorgrams were obtained by injection of the same volume of HBS-EP+ buffer in place of antibody Fab solution. Sensorgrams of the concentration series were corrected with corresponding blank curves and fitted globally with Biacore T200 evaluation software using a 1:1 Langmuir model of binding.

Affinity of antibody Fab to the His-tagged fusion peptide was measured on a Ni-NTA sensor chip (GE Healthcare). The Ni-NTA surface was activated by injection of 5 mM of Ni_2_SO_4_ in HBS-P+ buffer (10 mM HEPES, pH 7.4, 150 mM NaCl and 0.05% surfactant P-20) for 60 s at 6 μl/min and then stabilized by washing with HBS-EP+ buffer containing 3 mM EDTA for 60 s at 30 μl/min. Fusion peptide with His-tag at 20 ng/ml was captured at 6 μl/min flow rate for 60 s over the nickel activated sensor surface. Binding affinities for the 2G12-captured trimer were determined by using a serial dilution of antibody Fab solutions starting at 200nM during association phase. A dissociation phase at 30 μl/min for 300 s was used to determine binding kinetics. The surface was regenerated by flowing 300 mM imidazole to both channels at 6 μl/min for 60 s. Sensorgrams of the concentration series were corrected with corresponding blank curves and fitted globally with Biacore T200 evaluation software using a 1:1 Langmuir model of binding.

### HIV-1 Env mutagenesis

Site-directed mutagenesis on HIV-1 Env plasmids was performed through GeneImmune Biotechnology LLC, NY. T90A and S613A mutations were created to remove glycan 88 and 611, respectively.

### HIV-1 Env-pseudotyped virus

293T-grown HIV-1 Env-pseudotyped virus stocks were generated by cotransfection of the wildtype or mutant Env expression plasmids with a pSG3ΔEnv backbone^11^.

### Neutralization assays

A single round of entry neutralization assays using TZM-bl target cells were performed to assess monoclonal antibody neutralization as described^11^. Briefly, the monoclonal antibodies were tested via 5-fold serial dilutions starting at up to 500 μg/ml. Monoclonal antibodies were mixed with the virus stocks in a total volume of 50 μl and incubated at 37 °C for 1 hr. 20 μl of TZM-bl cells (0.5 million/ml) were then added to the mixture and incubated at 37 °C overnight. 130 μl cDMEM was added on day 2, and cells were lysed on day 3 and assessed for luciferase activity (RLU). The 50% and 80% inhibitory concentrations (IC_50_ and IC_80_) were determined using a hill slope regression analysis as described^11^.

To assess monoclonal antibody neutralization on a panel of 208 HIV-1 Env-pseudotyped viruses an automated 384-well microneutralization assay was performed as described previously^66^.

Serum neutralization was also assessed in the single round of entry neutralization assays using TZM-bl target cells, as described above. Before evaluation, all sera from immunized and control animal were heat-inactivated at 56 °C for 1 hr. All sera were tested via 4-fold serial dilutions starting at 1:20 dilution.

Serum neutralization with FP competition was performed with serum in presence of either PEGylated FP9 (AVGIGAVFL) or PEGylated non-cognate FLAG peptide. Mean and standard deviation (SD) of results from triplicated experiments were determined.

### Protein complex preparation

Antibody Fab and fusion peptide (residue 512-518) complexes were prepared by first dissolving fusion peptide in 100% DMSO at 50 mg/ml concentration, then mixing with Fab solution in 10:1 molar ratio to reach final protein complex centration of 15 mg/ml.

### Protein Crystal screening

Antibody Fab and fusion peptide (residue 512-518) complexes were screened for crystallization from JCSG1-4 protein crystal screening kits using a Cartesian Honeybee crystallization robot as described previously^15^ and a mosquito robot. Crystals initially observed from the wells were manually reproduced. vFP1.01/FP complex crystal grew in 0.2 M AmSO_4_, 0.1 M NaOAc pH 4.6; vFP7.04/FP complex crystal grew in 0.1 M MES pH 6.0, 30% PEG 6000; vFP16.02/FP complex crystal grew in 0.1 M NaOAc pH 4.5, 2 M AmSO_4_; vFP20.01/FP complex crystal grew in 0.1 M Citric acid pH 3.5, 2 M AmSO_4_; vFP5.01/FP complex crystals grew in 0.2 M MgCl_2_, 0.1 M Tris-HCl pH 8.5, 20% PEG 8000.

### X-ray data collection, structure solution, model building and refinement

Crystals were cryoprotected in 20% glycerol and flash-frozen in liquid nitrogen. Data were collected at a temperature of 100K and a wavelength of 1.00 Å at the SER-CAT beamline ID-22 (Advanced Photon Source, Argonne National Laboratory). Diffraction data were processed with the HKL2000 suite^67^. Structure solution was obtained by molecular replacement with Phaser using homologous Fab structures (PDB ID: 3BKY for vFP1-class antibody complex and 3LEY for vFP5.01 antibody complex) as search models. Model building was carried out with Coot^68^. Refinement was carried out with Phenix^69^. Ramachandran statistical analysis indicated that the final structures contained no disallowed or no more than 0.23% disallowed residues. Data collection and refinement statistics are shown in Supplementary Table 4.

### Cryo-EM data collection and processing

Env trimer used in cryo-EM was generated in GNTI-cell line as described previously^41^. To prepare Env complexes, BG505 DS-SOSIP at a final concentration of 0.3-0.5 mg/ml was incubated with 4-5-fold molar excess of the antibody Fab fragments for 30-60 minutes. To prevent aggregation during vitrification, the sample was incubated in 0.085 mM dodecyl-maltoside (DDM). The vFP1.01 and vFP5.01 bound complexes were vitrified by applying 3 μl of sample to freshly plasma-cleaned C-flat holey carbon grids (CF-1.2/1.3-4C) (EMS, Hatfield, PA) for vFP1.01 and gold grids for vFP5.01, allowing the sample to adsorb to the grid for 60 s, followed by blotting with filter paper and plunge-freezing into liquid ethane using the CP3 cryo-plunger (Gatan, Inc.) (20°C, 85-90% relative humidity).

The vFP16.02 and vFP20.01 bound complexes were vitrified using a semi-automated Spotiton V1.0 robot^70^ The grids used were specially designed Nanowire self-blotting grids with a Carbon Lacey supporting substrate. Sample was dispensed onto these nanowire grids using a picoliter piezo dispensing head. A total of ∼5 nl sample was dispensed in a stripe across each grid, followed by a pause of a few milliseconds, before the grid was plunged into liquid ethane.

Data were acquired using the Leginon system installed on Titan Krios electron microscopes operating at 300kV and fitted with Gatan K2 Summit direct detection device. The dose was fractionated over 50 raw frames and collected over a 10 s exposure time. Individual frames were aligned and dose-weighted.

CTF was estimated using the GCTF package^71^. Particles were picked using DoG Picker within the Appion pipeline^72^. 2D and 3D classifications were performed using RELION^73^. A map of unliganded BG505 SOSIP.664 (EMDB ID 5782), low-pass filtered to 60 Å was used as the starting point of 3D classification followed by 3D refinement in either RELION or cryoSparc^74^. For the vFP16.02 and vFP20.01 complexes, after 3D classification in RELION, an additional step of *ab initio* reconstruction was performed using cryoSparc.

### Cryo-EM Model fitting

Fits of HIV-1 trimer and Fab to the cryo-EM reconstructed maps were performed using Chimera^75^. Glycosylated BG505 SOSIP trimer structure (PDB ID: 5YFL) was used for the trimer fits. For antibody fitting, we used the fusion peptide-bound coordinates of vFP1.01 and vFP5.01. For the antibody Fabs, both orientations rotated ∼180° about the Fab longitudinal axis were tested, and the optimal fit was decided based on map-to-model correlation and positioning of the fusion peptide bound to the Fab relative to Env. For the vFP16.02 and vFP20.01 bound complexes, the coordinates were further fit to the electron density by an iterative process of manual fitting using Coot^68^ and real space refinement within Phenix^69^. Molprobity^76^ and EMRinger^77^ were used to check geometry and evaluate structures at each iteration step. Figures were generated in UCSF Chimera and PyMOL. Map-fitting cross correlations were calculated using Fit-in-Map feature in UCSF Chimera. Map-to-model FSC curves were generated using EMAN2. Local resolution of cryo-EM maps was determined using RELION.

### Defining vFP1 class antibody V-gene sequence signature

The V-gene sequence signature for vFP1 class antibodies were defined by examining the vFP1 class antibody sequences that neutralize at least seven out of the ten tested isolates and the structures of FP in complex with vFP1.01, vFP16.02, and vFP20.01. A residue position was considered as part of the sequence signature if at least one side chain heavy atom was within five angstroms from any fusion peptide heavy atom for all three complex structures, and no more than three similar amino acid types have a combined prevalence of more than 90%, and each of these amino acid types had a prevalence of more than 10%.

### Molecular dynamics simulation of mannose-5 Env trimer

By using the BG505 SOSIP.664 Env trimer structure (PDB ID: 4TVP) as a starting template, we modeled in a fully extended mannose 5 moiety at each *N*-linked glycosylation sequon. The fusion peptide structure was then grafted onto our full mannose 5 model followed by 5000 steps of conjugate gradient energy minimization in implicit solvent using NAMD. The obtained structure was then solvated in a 17 Å padding water box, neutralized by the addition of NaCl at a concentration of 150 mM. The CHARMM force field (https://www.charmm.org) was used for the parameterization of the protein (including CMAP corrections and the mannose 9). TIP3P water parameterization^78^ was used to describe the water molecules.

Two independent molecular simulation were carried out using ACEMD molecular dynamics software on their METROCUBO workstation (https://www.acellera.com/products/GPU-hardware-molecular-dynamics-metrocubo/). The system was minimized for 2000 steps, followed by equilibration using the NPT ensemble for 50 ns at 1 atm and 300 K using a time-step of 2 fs. We also used rigid bonds and a 9 Å cutoff using PME for long range electrostatics. During the equilibration phase, heavy atoms on the protein were constrained by a 1 kcal/molÅ-2 spring constant and slowly relaxed over the first 5 ns. Following the relaxation phase, the protein was allowed to move freely and simulated for 500 ns under the NVT ensemble using ACEMD’s NVT ensemble with a Langevin thermostat. To achieve a time-step of 4 ps, we used damping at 0.1 ps-1 and a hydrogen mass repartitioning scheme. Each simulation ran up to 500 ns.

The conformations of the fusion peptide (residue 512-519) were extracted from the MD simulations every 100 ps, producing an ensemble of 30’000 structures. Prody (http://prody.csb.pitt.edu) was used to perform the principal component analysis of backbone atoms. The conformations of five crystalized fusion peptides were then projected into the eigenspace defined by the first two components: vFP1.01, vF5.01, PGT151 (PDB: 5FUU), VRC34.01 (PDB: 5I8H) and clade G (PDB: 5FYJ).

### Analysis of antibody angle of approach to HIV-1 Env

To compare modes of antibody recognition of HIV-1 Env by vaccine elicited fusion peptide antibodies and VRC34.01, structural models of antibody in complex with HIV-1 BG505 SOSIP Env trimer derived from x-ray crystallography and EM were superimposed by pairwise alignment of the Cα atom coordinates of the HIV-1 Env. To simplify the comparison, we first defined two common axes on the HIV-1 Env, the trimer axis and the protomer axis, as reference lines. The trimer axis was defined by two points, each with x, y, z coordinates obtained by averaging the coordinates of the Cα atom of a residue and its 3-fold symmetry mates on the same trimer. The protomer axis was defined by a line perpendicular to the trimer axis that passes the center of the protomer. On the antibody side, we also defined two axes for each Fab. The long axis of a Fab was defined by two points, one point from the variable domain with x, y, z coordinates obtained by averaging the coordinates of the Cα atom of the 4 conserved Cys (Cys 22 and Cys92 of heavy chain, and Cys23 and Cys88 of light chain), and the other from the constant domain with x, y, z coordinates obtained by averaging the coordinates of the Cα atom of the 4 conserved Cys (Cys 140 and Cys196 of heavy chain, and Cys134 and Cys194 of light chain). The short axis of a Fab was defined by a line connecting Cα atoms of heavy chain Cys22 and light chain Cys23. Approaching angles of each antibody were then calculated as 1) angle between trimer axis and Fab long axis, and 2) angle between the axis of major interaction protomer and Fab long axis. In addition, the relative heavy and light chain orientation of antibody variable domains was compared by angles between the Fab short axes. The axes were visualized in PyMOL by placing their coordinates in PDB format.

### Autoreactivity assay

Antibodies were assessed for autoreactivity by testing for binding to HEp2 cells by indirect immunofluorescence (Zeus Scientific, ANA HEp2 test system) and cardiolipin by ELISA (Inova Diagnostics, QUANTA Lite ACA IgG III), per the manufacturer’s instructions. On HEp2 cells, antibodies were assigned a score between 0 and 3+ using control antibodies as reference. In the cardiolipin binding assay, OD values were converted to GPLs using standard samples provided in the kit. Monoclonal antibodies that scored greater than 20 GPLs at 33 μg/ml were considered autoreactive.

### Neutralization fingerprinting analysis

The neutralization fingerprint of a monoclonal antibody or polyclonal plasma is defined as the potency pattern with which the antibody/plasma neutralizes a set of diverse viral strains. The neutralization fingerprints of a set of monoclonal antibodies and NHP plasma were compared and clustered according to fingerprint similarity, as described previously^79^. A set of 132 strains^80^ was used in the neutralization fingerprint analysis for Fig. S11, and the 58 FP-selected strains for Fig. 6.

### Associations between glycosylation patterns versus neutralization

We calculated the associations between the sequence variability of FP neighboring glycosylation sites (HXB2 numbering 88, 241, 448 and 611) and the large panel neutralization using an approach implemented with R package SeqFeatR (https://seqfeatr.zmb.uni-due.de). Briefly, for each residue position we create contingency table based on glycan occurrence and neutralization sensitivity and evaluate the association using Fisher’s exact test. The resulting P-values were corrected using multiple testing method Holm.

### Statistical analysis

The correlation between neutralization breadth on 58-isolate panel with FP8=AVGIGAVF and neutralization breadth on the full 208-isolate panel was performed using Pearson correlation. The differences of number of neutralized ioslates between FP-KLH primed+DS-SOSIP boosted group and DS-SOSIP alone group were evaluated using one-tailed Mann-Whitney test. The differences of neutralization betwewen FP-KLH groups and KLH-only group were evaluated using one-tailed Mann-Whitney test. The correlation between antibody trimer binding and antibody neutralization breadth was performed using Pearson correlation. The differences of divergence of germline between antibodies from trimer-primed groups and antibodies from non-trimer-primed groups were evaluated using two-tailed Mann-Whtiney test. The differences of sera neutralization between trimer-boosted animals and non-trimer-boosted animals were evaluated using one-tailed Mann-Whitney test. The association of FP-proximal N-glycan sequons (N88, N241, N448, and N611) and neutralization based on the 208-isolate panel was evaluated using two-tailed Fisher’s exact test. Unless otherwise indicated, all data were plotted and graphed using GraphPad Prism, version 7.0. P < 0.05 indicated a statistically significant difference.

### Code availability

No studies deemed central to the conclusions were carried out with custom code.

### Data availability

All data generated or analyzed during this study are included in this published article (and its Supplementary Information files). Atomic coordinates and structure factors of the reported crystal structures were deposited in the Protein Data Bank with PDB ID 5TKJ, 5TKK, 6CDM, 6CDO, 6CDP. Cryo-EM reconstructions were deposited in the Electron Microscopy Data Bank with EMDB accession code EMD-7460, EMD-7459, EMD-8420, EMD-8421 and EMD-8422, and in Protein Data Bank with PDB ID 6CDI and 6CDE. Heavy chain- and light chain-variable sequences of monoclonal antibodies vFP1.01 - vFP7.05 and vFP7.06 - vFP 32.07 were deposited with GenBank under accession numbers KX949064 - KX949087 and MH017667 - MH017826, respectively.

### Life Sciences Reporting Summary

Further information on experimental design is available in the Life Sciences Reporting Summary.

